# Salt-inducible kinases (SIKs) regulate TGFβ-mediated transcriptional and apoptotic responses

**DOI:** 10.1101/809228

**Authors:** Luke D. Hutchinson, Nicola J. Darling, Stephanos Nicolaou, Ilaria Gori, Daniel R. Squair, Philip Cohen, Caroline S. Hill, Gopal P. Sapkota

**Affiliations:** MRC Protein Phosphorylation and Ubiquitylation Unit, School of Life Sciences, University of Dundee, Sir James Black Centre, Dow Street, Dundee, DD1 5EH, United Kingdom; The Francis Crick Institute, 1 Midland Road, London, NW1 1AT, United Kingdom

## Abstract

The signalling pathways initiated by members of the transforming growth factor-β (TGFβ) family of cytokines control many metazoan cellular processes, including proliferation and differentiation, epithelial-mesenchymal transition (EMT), and apoptosis. TGFβ signalling is therefore strictly regulated to ensure appropriate context-dependent physiological responses. In an attempt to identify novel regulatory components of the TGFβ signalling pathway, we performed a pharmacological screen using a cell line engineered to report the endogenous transcription of the TGFβ-responsive target gene *PAI-1*. The screen revealed that small-molecular inhibitors of salt-inducible kinases (SIKs) attenuate TGFβ-mediated transcription of *PAI-1* without affecting receptor-mediated SMAD phosphorylation, SMAD complex formation or nuclear translocation. We provide evidence that genetic inactivation of SIK isoforms also attenuates TGFβ-dependent transcriptional responses. Pharmacological inhibition of SIKs using multiple small-molecule inhibitors potentiated apoptotic cell death induced by TGFβ stimulation. Our data therefore provides evidence for a novel function of SIKs in modulating TGFβ-mediated transcriptional and cellular responses.

## Introduction

Signalling pathways initiated by the TGFβ family of cytokines are amongst the most prevalent and diverse in metazoan biology and regulate a multitude of processes including cellular proliferation and differentiation, epithelial-mesenchymal transition (EMT), cell migration, immunoregulation and apoptotic cell death in a context-dependent manner^1–6^. Consequently, perturbations within the signalling pathway have been associated with the pathogenesis of many human disorders including cancer. For example, in normal epithelial cells, TGFβ has a tumour suppressive function, principally through its ability to induce cytostasis and apoptotic cell death^7–9^. In contrast, during tumour progression, the suppressive effect of TGFβ is lost and, in certain cancers, corruption of the signalling pathway can result in TGFβ exerting a pro-oncogenic effect^7, 10, 11^. Inhibition of the TGFβ pathway has therefore been proposed as a potential therapeutic strategy in certain pathological contexts^12, 13^. However, the highly pleiotropic and context-dependent nature of TGFβ signalling has provided a considerable challenge for pharmacological intervention^14^. Elucidating the context-dependent regulatory mechanisms underlying TGFβ signalling is therefore of considerable importance in identifying novel therapeutic interventions.

TGFβ signalling is initiated upon the binding of TGFβ ligand dimers to cognate transmembrane receptor serine-threonine protein kinases to form activated heterotetrameric receptor complexes containing two type I receptors and two type II receptors^15^. This allows the constitutively active type II receptor to phosphorylate multiple serine and threonine residues within the cytoplasmic domain of the type I receptor, which enables the type I receptor to bind and phosphorylate the SMAD transcription factors 2/3 (SMADs2/3) at the Ser-Xxx-Ser motif at the carboxy-terminal tail^16–18^. Receptor-mediated phosphorylation of R-SMADs facilitates interaction with the co-SMAD, SMAD4, followed by accumulation in the nucleus, where the SMAD complex co-operates with different transcriptional co-regulators to modulate the expression of hundreds of target genes in a cell type- and context-dependent manner^18–20^.

Previously, we developed an endogenous transcriptional reporter cell line for the TGFβ pathway using CRISPR-Cas9 genome editing technology^21^ by inserting firefly (*Photinus pyralis*) luciferase and green fluorescent protein (GFP) at the native TGFβ-responsive target gene *plasminogen activator inhibitor 1* (*PAI-1*) locus (Figure 1A). The transcription of *PAI-1* is induced in response to TGFβ signals in different cell types in a SMAD-dependent manner^22, 23^. Moreover, the promoter region of the endogenous *PAI-1* gene has been frequently utilised in order to generate conventional luciferase-based overexpression reporter systems for the study of TGFβ-mediated transcriptional regulation^24^. In order to identify novel regulatory components of the TGFβ pathway, we performed a pharmacological screen in this endogenous TGFβ-responsive transcriptional reporter cell line using a panel of small-molecules obtained from the MRC International Centre for Kinase Profiling at the University of Dundee. The panel consisted predominantly of selective and potent inhibitors of protein kinases but also included a small number of compounds which target components of the ubiquitin-proteasome system (UPS). The screen identified salt-inducible kinases (SIKs), which are members of the AMPK-activated protein kinase (AMPK)-related subfamily of serine-threonine specific kinases^25, 26^, as potential novel regulators of TGFβ-mediated gene transcription. In this study, we have therefore investigated the role of SIKs in regulating the TGFβ signalling pathway.

**Figure 1.**
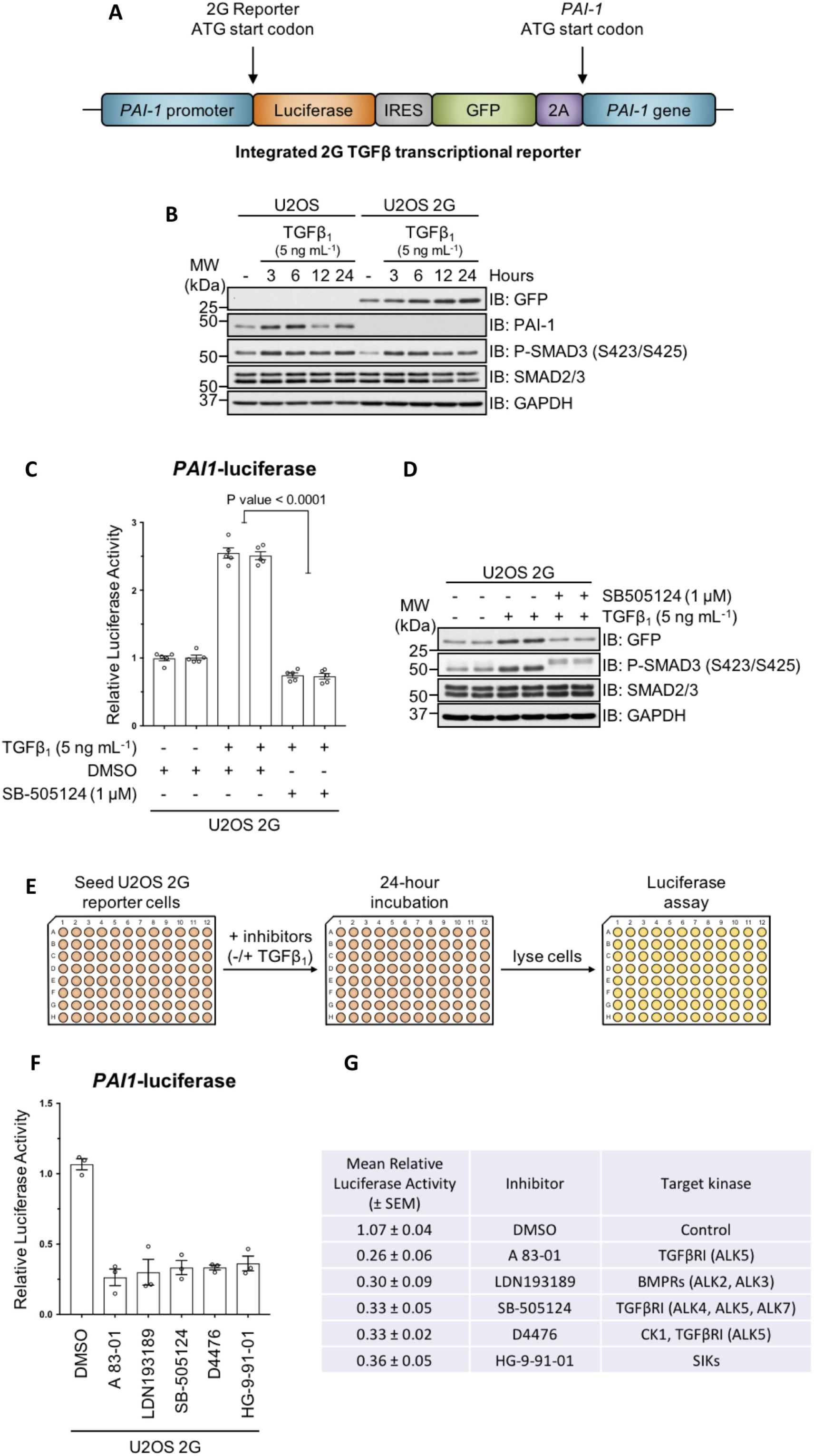
Pharmacological screen in endogenous TGFβ transcriptional reporter cells. **A:** Schematic representation of the dual-reporter cassette inserted in-frame with the ATG start codon of the endogenous *PAI-1* gene in U2OS human osteosarcoma cells. **B:** Immunoblot analysis of wild type U2OS and U2OS 2G transcriptional reporter cell lines stimulated with TGFβ_1_ (5 ng mL^-1^) for the indicated durations. Cell lysates were resolved via SDS-PAGE and membranes subjected to immunoblotting with the indicated antibodies. **C:** Luciferase assay analysis of U2OS 2G transcriptional reporter cells incubated with either SB-505124 or DMSO control in the presence of TGFβ_1_ stimulation. **D:** Immunoblot analysis of U2OS transcriptional reporter cells incubated with either SB-505124 or DMSO control in the presence of TGFβ_1_ stimulation. Cell lysates were resolved via SDS-PAGE and membranes subjected to immunoblotting with the indicated antibodies. **E:** Schematic representation of the experimental workflow for the pharmacological screen in U2OS 2G transcriptional reporter cells. **F and G:** The top 5 hits obtained from three independent experiments which reduced TGFβ-induced luciferase activity. Data indicates the mean luciferase activity values (± SEM) relative to internal DMSO controls.

## Results

### Identification of salt-inducible kinases (SIKs) as novel regulators of TGFβ-mediated gene transcription

We tested the utility of the endogenous TGFβ-responsive transcriptional reporter U2OS cell line (U2OS-2G) (Figure 1A) for a pharmacological screen. Stimulation of wild type (WT) U2OS and U2OS-2G cells with TGFβ_1_ over 24 hours resulted in time-dependent induction of PAI-1 and GFP expression respectively (Figure 1B), and comparable levels of SMAD3 *C*-terminal phosphorylation in both cell lines. TGFβ induced a significant increase in relative luciferase activity in U2OS-2G cells over unstimulated cells, which was blocked with SB-505124, a selective inhibitor of the TGFβ type I receptor (TGFβR1) kinases^27, 28^ (Figure 1C). Similarly, TGFβ-induced GFP expression in U2OS-2G cells is blocked with SB-505124 (Figure 1D). These data confirmed the suitability of U2OS-2G cells for pharmacological screens. A 96-well plate format pharmacological screen was performed to identify potential novel regulators of the TGFβ pathway (Figure 1E). TGFβR1 inhibitors SB-505124^27, 28^ and A 83-01^29^ served as positive controls, while DMSO a negative control. All compounds were used at 1 µM final concentration. Both SB-505124 and A 83-01 significantly inhibited TGFβ-induced luciferase activity compared with DMSO controls (Figure 1F and 1G). Additionally, D4476 and LDN193189, also significantly inhibited TGFβ-induced luciferase activity. D4476 was initially identified as an inhibitor of TGFβR1^30^, although subsequent *in vitro* profiling revealed that is inhibited casein kinase 1 (CK1) with greater potency^31^. LDN193189 is an ATP-competitive inhibitor of the BMP type I receptor kinases^32^. However, at 1 µM concentration it also inhibits the TGFβR1^28^. The majority of the compounds used in the screen did not significantly affect TGFβ-induced luciferase reporter activity. Interestingly however, we observed that HG-9-91-01, a potent ATP-competitive inhibitor of salt-inducible kinase (SIK) isoforms^33^, significantly attenuated TGFβ-induced luciferase activity (Figure 1F and 1G), suggesting a possible role for SIKs in TGFβ-induced transcription.

### Characterisation of SIK inhibitors in the context of TGFβ signalling

To explore the role of SIKs in TGFβ signalling further, in addition to HG-9-91-01, we utilised MRT199665, a structurally distinct inhibitor of SIK isoforms^33^ (Figure 2A). MRT199665 also suppressed TGFβ-induced luciferase activity in U2OS-2G cells, as potently as SB-505124 and HG-9-91-01 (Figure 2B). Both HG-9-91-01 and MRT199665 inhibited the phosphorylation of a known SIK substrate CRTC3 at S370^33–35^ compared with DMSO control (Figure 2C). Because kinase inhibitors often display off-target inhibition, we tested whether the attenuation of TGFβ-induced luciferase activity by HG-9-91-01 and MRT199665 occurred as a result of the off-target inhibition of the TGFβR1 upstream of SMAD2/SMAD3 phosphorylation. HG-9-91-01 substantially inhibited TGFβ-induced SMAD3 phosphorylation, to a similar extent as SB-505124, compared to DMSO controls, whereas MRT199665 did not (Figure 2D), suggesting that HG-9-91-01 could inhibit either type I or type II TGFβ receptors. Indeed, at concentrations of 0.1, 1 and 10 µM *in vitro*, HG-9-91-01 inhibited TGFβR1 (ALK5) kinase activity, whereas MRT199664 did not (Figure 2E). Because of this off-target inhibition of TGFβR1 by HG-9-91-01, we decided to employ MRT199665 as SIK inhibitor for subsequent experiments.

**Figure 2.**
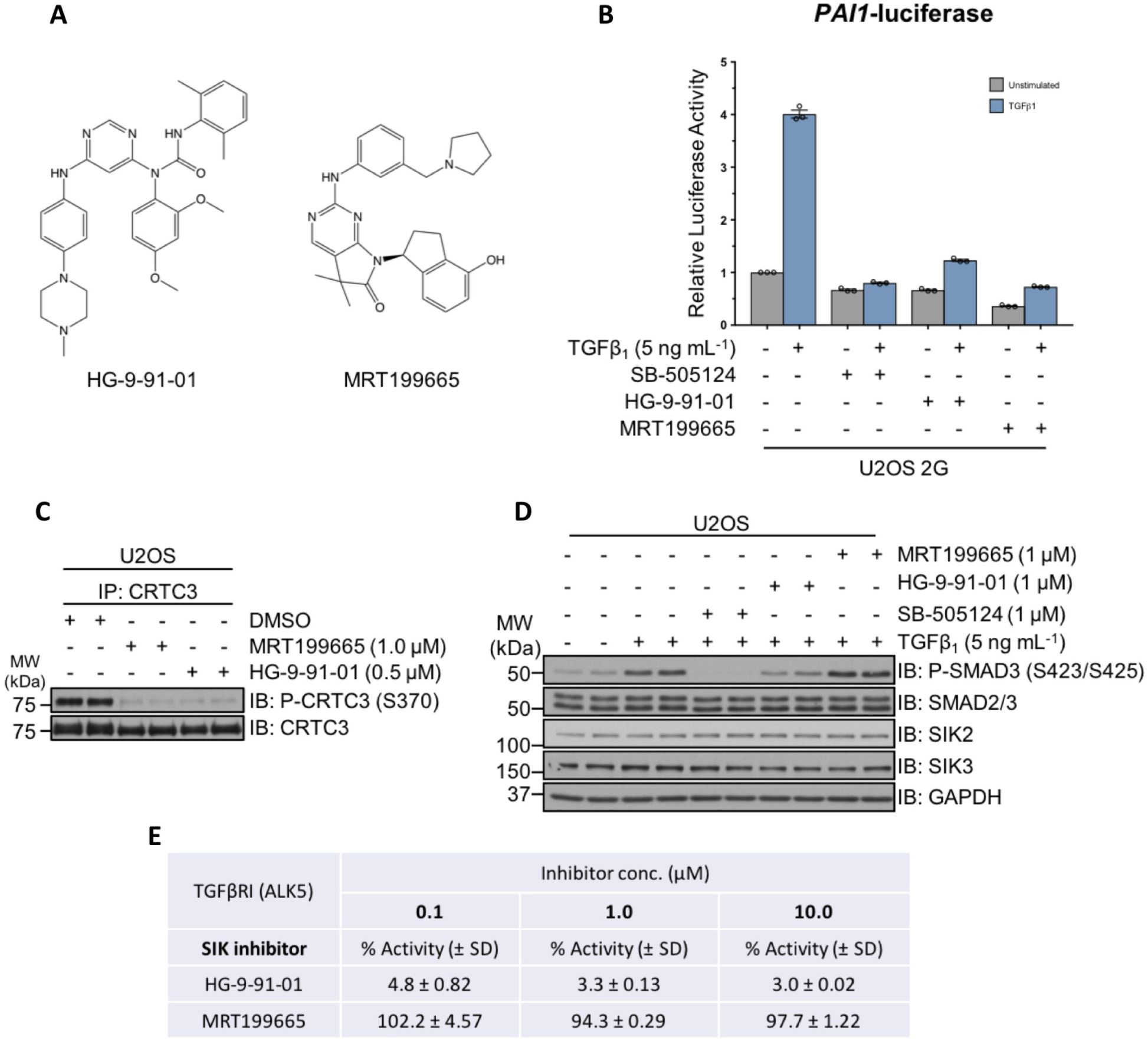
Characterisation of pharmacological SIK inhibitors in the context of TGFβ signalling. **A:** The chemical structures of HG-9-91-01 and MRT199665, ATP-competitive small-molecule inhibitors of SIK isoforms. **B:** Luciferase assay analysis of U2OS 2G transcriptional reporter cells incubated with either DMSO, SB-505124, HG-9-91-01 or MRT199665, in the presence or absence of TGFβ_1_ stimulation. **C:** Immunoblot analysis of endogenous CRTC3 phosphorylation in wild type U2OS cells following incubation with DMSO, MRT199665 or HG-9-91-01. Cell lysates were subjected to endogenous CRTC3 immunoprecipitation (IP) and subsequently resolved via SDS-PAGE. Membranes were subjected to immunoblotting with the indicated antibodies. **D:** Immunoblot analysis of wild type U2OS cells incubated with either SB-505124, HG-9-91-01 or MRT199665 in the presence of TGFβ_1_ stimulation. Cell lysates were resolved via SDS-PAGE and membranes subjected to immunoblotting with the indicated antibodies. **E:** *In vitro* kinase assay analysis of recombinant constitutively active TGFβR1 (ALK5) in the presence of HG-9-91-01 or MRT199665 at the three indicated concentrations. Values denote the mean percentage activity remaining (± SD).

### MRT199665 attenuates the expression of endogenous TGFβ target genes

Any compound that inhibits luciferase enzymatic activity could potentially yield a false-positive result in U2OS-2G cells. Therefore, to exclude this possibility for MRT199665, we tested whether the TGFβ-induced GFP expression in U2OS-2G cells was affected by MRT199665. The TGFβ-induced expression of GFP in U2OS-2G cells was inhibited with MRT199665, to the same extent as SB-505124, compared to DMSO control (Figure 3A), while the TGFβ-induced SMAD2/SMAD3 *C*-terminal phosphorylation was unaffected by MRT199665 (Figure 3A). Moreover, MRT199665 significantly attenuated TGFβ-induced *PAI-1* mRNA expression in WT U2OS cells (Figure 3B). In WT A-172 human glioblastoma cells, MRT199665 also inhibited TGFβ-induced expression of *PAI-1* mRNA, as well *SMAD7* and connective tissue growth factor (*CTGF*) mRNAs (Figure 3C). These data demonstrate that MRT199665 inhibits TGFβ-induced transcription of endogenous target genes in different cells without affecting SMAD2/SMAD3 phosphorylation.

**Figure 3.**
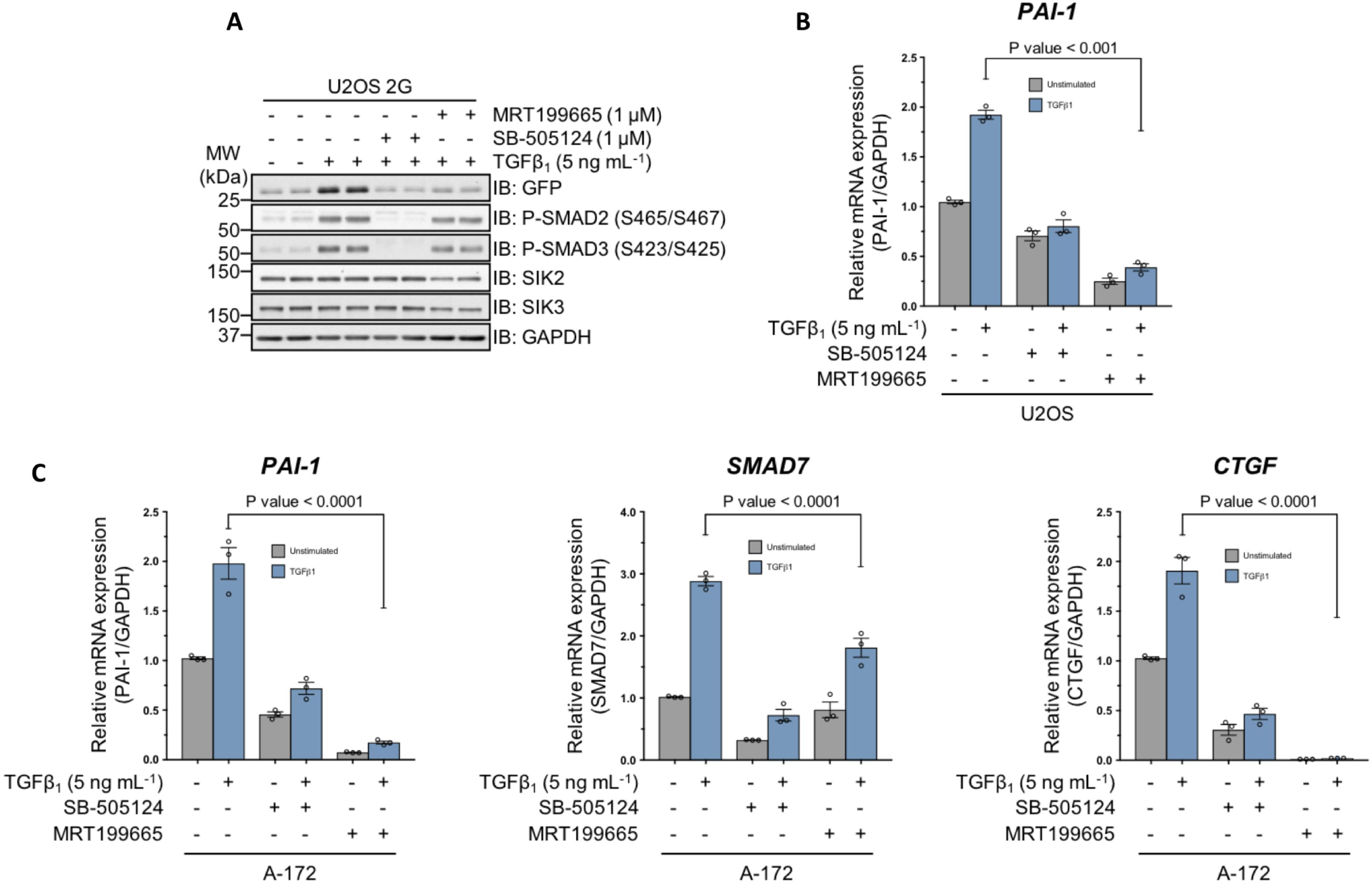
MRT199665 attenuates TGFβ-mediated transcription in human cancer cell lines A: Immunoblot analysis of U2OS 2G transcriptional reporter cells incubated with either SB-505124 or MRT199665 in the presence of TGFβ_1_ stimulation. Cell lysates were resolved via SDS-PAGE and membranes subjected to immunoblotting with the indicated antibodies. **B:** RT-qPCR analysis of *PAI-1* mRNA expression in wild type U2OS human osteosarcoma cells incubated with either SB-505124 or MRT199665 in the presence or absence of TGFβ_1_ stimulation. **C:** RT-qPCR analysis of *PAI-1*, *SMAD7* and *CTGF* mRNA expression in wild type A-172 human glioblastoma cells incubated with either SB-505124 or MRT199665 in the presence or absence of TGFβ_1_ stimulation.

### Genetic inactivation of SIK2/3 attenuates the TGFβ-mediated induction of PAI-1 expression

We employed genetic approaches to test the impact of SIK kinase activity on TGFβ signalling. SIKs are members of the AMP-activated protein kinase (AMPK)-related subfamily of serine-threonine protein kinases that require LKB1-mediated phosphorylation of a conserved threonine residue within the activation loop in order to become catalytically active^25, 26^ (Figure 4A). In LKB1-deficient WT HeLa cells^36–38^, TGFβ_1_ induced a 1.5-fold increase in *PAI-1* mRNA expression relative to unstimulated controls. However, stable overexpression of catalytically active LKB1 (LKB1^WT^), but not catalytically inactive mutant (LKB1^D194A^), in WT HeLa cells significantly enhanced the TGFβ-induced transcription of *PAI-1* mRNA (Figure 4B) as well as PAI-1 protein levels (Figure 4C), although the levels of LKB1^WT^ restored in HeLa cells was substantially higher than the LKB1^D194A^ mutant (Figure 4C).

**Figure 4.**
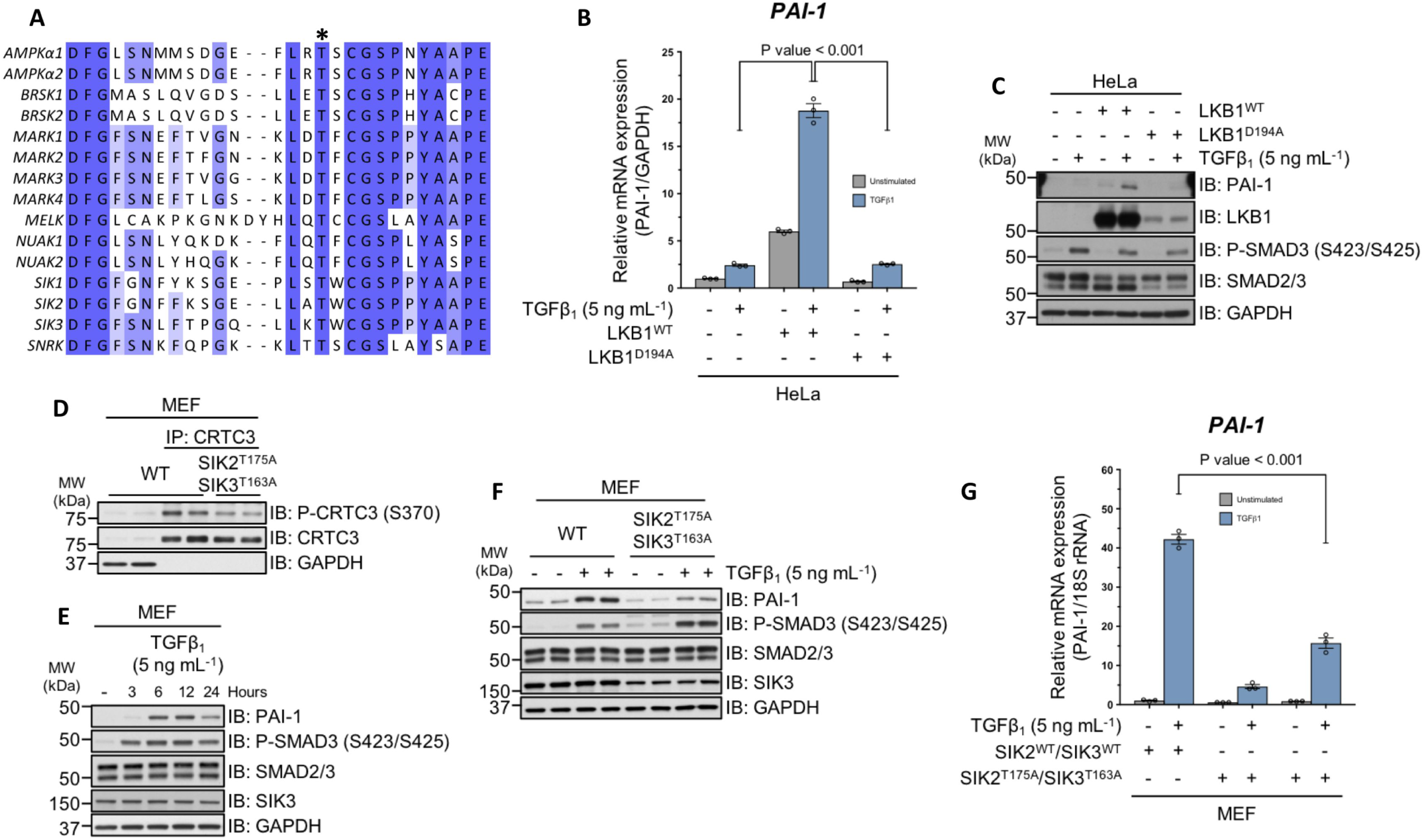
Genetic evidence for the involvement of SIK isoforms in TGFβ-mediated PAI-1 expression. **A:** Sequence alignment of the activation segment of the human AMPKα catalytic subunits and the 13 members of the AMPK-related family of protein kinases. The asterisk indicates the conserved activation (T) loop threonine residue that is phosphorylated by LKB1. **B:** RT-qPCR analysis of *PAI-1* mRNA expression in wild type HeLa cervical adenocarcinoma cells and HeLa cells overexpressing either LKB1^WT^ or LKB1^D194A^ following TGFβ_1_ stimulation. **C:** Immunoblot analysis of wild type HeLa cells and HeLa cells overexpressing either LKB1^WT^ or LKB1^D194A^ following TGFβ_1_ stimulation. Cell lysates were resolved via SDS-PAGE and membranes subjected to immunoblotting with the indicated antibodies. **D:** Immunoblot analysis of endogenous CRTC3 phosphorylation in wild type MEFs and MEFs derived from homozygous SIK2^T175A^/SIK3^T163A^ KI mouse embryos. Cell lysates were subjected to endogenous CRTC3 IP and subsequently resolved via SDS-PAGE. Membranes were subjected to immunoblotting with the indicated antibodies. **E:** Immunoblot analysis of wild type MEFs stimulated with TGFβ_1_ for the indicated durations. Cell lysates were resolved via SDS-PAGE and membranes subjected to immunoblotting with the indicated antibodies. **F:** Immunoblot analysis of wild type and homozygous SIK2^T175A^/SIK3^T163A^ MEFs. Cell lysates were resolved via SDS-PAGE and membranes subjected to immunoblotting with the indicated antibodies. **G:** RT-qPCR analysis of *PAI-1* mRNA expression in wild type and homozygous SIK2^T175A^/SIK3^T163A^ KI MEFs following TGFβ_1_ stimulation.

The catalytic activity of SIK isoforms can be ablated via mutation of the activation loop threonine to alanine^39^, which abolishes LKB1-mediated phosphorylation. Indeed, in mouse embryonic fibroblasts (MEFs) derived from embryos harbouring homozygous SIK2^T175A^ and SIK3^T163A^ genotypes^39^, the phosphorylation of CRTC3 at S370 is substantially reduced compared to WT control MEFs (Figure 4D). A time-course treatment WT MEFs with TGFβ_1_ resulted in robust SMAD3 phosphorylation and increase in PAI-1 protein levels at 6 hours (Figure 4E). When homozygous SIK2^T175A^/SIK3^T163A^ MEFs were subjected to TGFβ stimulation for 6 hours, the induction of PAI-1 protein expression was substantially attenuated compared with WT MEFs, despite the observation of higher SMAD3 phosphorylation in the SIK2^T175A^/SIK3^T163A^ mutant MEFs (Figure 4F). Consistent with this, the relative *PAI-1* mRNA expression in response to TGFβ stimulation was significantly reduced in MEFs derived from two independent homozygous SIK2^T175A^/SIK3^T163A^ mice relative to WT MEFs (Figure 4G).

### Impact of SIK isoforms on TGFβ-dependent proliferative responses

TGFβ inhibits epithelial cell proliferation, in part through transcriptional upregulation of cyclin-dependent kinase (CDK) inhibitors p21^CIP1^ and p27^KIP1^ and downregulation of the proto-oncogene c-Myc^1,4, 7^. In HaCaT cells, we observed an increase in endogenous p27^KIP1^ and p21^CIP1^ protein levels and a decrease in c-Myc protein levels over a 24-hour time course of TGFβ treatment (Figure 5A). Rather surprisingly, treatment of HaCaT cells with MRT199665 resulted in increased expression of both p21^CIP1^ and p27^KIP1^ even in the absence of TGFβ treatment, and this increase was more pronounced after stimulation of cells with TGFβ compared to DMSO controls (Figure 5B). This suggested that inhibition of SIKs alone may exert cytostatic effects. When we analysed the proliferation of HaCaT cells over a period of 160 hours, the control cells displayed a typical sigmoid growth curve, while TGFβ treatment caused a significant inhibition of proliferation after 100 hours (Figure 5C). Under these conditions, MRT199665 profoundly suppressed cell proliferation at all time points, regardless of TGFβ treatment (Figure 5C). We also observed a substantial increase in p27^KIP1^ levels in MEFs derived from homozygous SIK2^T175A^/SIK3^T163A^ KI mice relative to WT mice irrespective of TGFβ stimulation (Figure 5D), suggesting that SIK kinase activity plays a fundamental role in suppressing p27^KIP1^ protein levels. To further confirm that SIK inhibition promotes cytostasis independent of TGFβ stimulation, we exploited SMAD3^-/-^ HaCaT cells and showed that treatment of both WT and SMAD3^-/-^ HaCaT cells with MRT199665 resulted in an increase in p21^CIP1^ and p27^KIP1^ levels compared with untreated control cells (Figure 5E).

**Figure 5.**
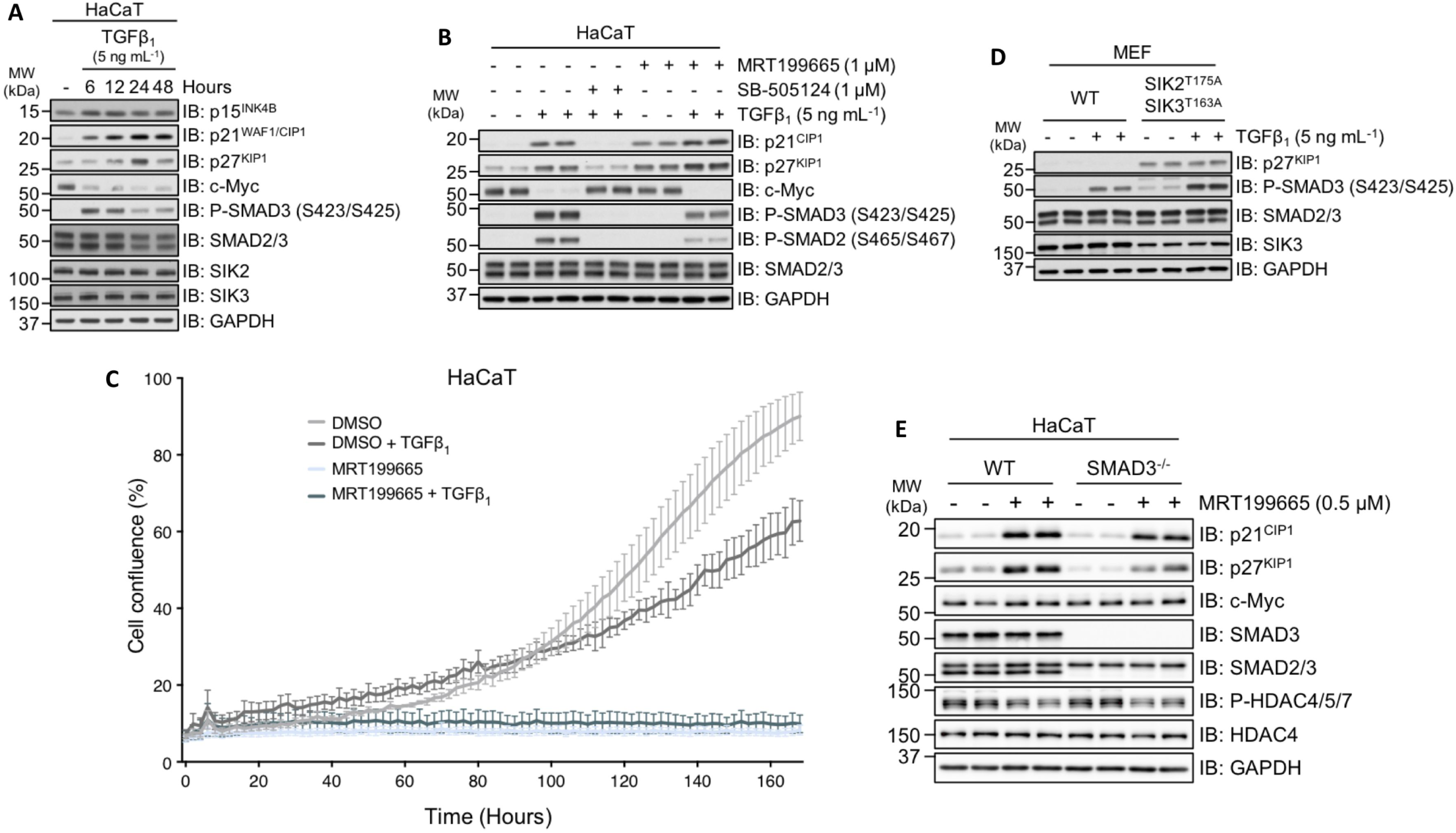
SIK inhibition enhances TGFβ-independent cytostasis. **A:** Immunoblot analysis of wild type HaCaT cells stimulated with TGFβ_1_ for the indicated durations. Cell lysates were resolved via SDS-PAGE and membranes subjected to immunoblotting with the indicated antibodies. **B:** Immunoblot analysis of wild type HaCaT cells incubated with either DMSO, SB-505124 or MRT199665, in the presence or absence of TGFβ_1_ stimulation. Cell lysates were resolved via SDS-PAGE and membranes subjected to immunoblotting with the indicated antibodies. **C:** Immunoblot analysis of wild type and homozygous SIK2^T175A^/SIK3^T163A^ MEFs following TGFβ_1_ stimulation. Cell lysates were resolved via SDS-PAGE and membranes subjected to immunoblotting with the indicated antibodies. **D:** Immunoblot analysis of wild type and SMAD3^-/-^ HaCaT cells incubated in the presence or absence of MRT199665. Cell lysates were resolved via SDS-PAGE and membranes subjected to immunoblotting with the indicated antibodies.

In many epithelial cells TGFβ-induced cytostasis, through the induction of p21^CIP1^ and p27^KIP1^ and suppression of c-Myc, is often necessary for subsequent TGFβ-dependent cell fates, such as differentiation, epithelial-mesenchymal transition (EMT) and apoptosis. Our unexpected findings that inhibition of SIK isoforms induces the expression of p21^CIP1^ and p27^KIP1^ levels independent of TGFβ prompted us to explore whether TGFβ-induced epithelial cell fates are sensitised by SIK inhibitors. We therefore sought to investigate whether inhibition of SIK isoforms sensitises cells to TGFβ-induced EMT and apoptosis.

### Pharmacological inhibition of SIKs potentiates TGFβ-mediated apoptosis

NMuMG murine mammary epithelial cells undergo both EMT and apoptosis upon TGFβ stimulation^40–44^. When we tested the effect of MRT199665 on TGFβ-induced EMT in NMuMG cells, which usually takes around 24-48 hours, it became apparent that there was profound cell death within 12-24 hours, prompting us to investigate apoptosis. The apoptotic response to TGFβ is mediated in part by the executioner cysteine-aspartic acid protease, caspase-3. Caspase-3 is synthesised as an inactive proenzyme and requires proteolytic cleavage in order to become catalytically active. Apoptosis can thus be monitored via the detection of the cleaved, and hence activated, form of caspase-3. Furthermore, activated caspase-3 mediates the proteolytic cleavage of poly (ADP-ribose) polymerase (PARP), which can also be used to monitor apoptosis. Stimulation of NMuMG cells with TGFβ_1_ over a period of 72 hours induced the activation of caspase-3 and subsequent cleavage of PARP, with maximal cleavage observed at 24 hours (Figure 6A). The expression of pro-apoptotic factor Bim was observed at 48-72 hours (Figure 6A). In these cells, SIK inhibitors HG-9-91-01 and MRT199665 resulted in reduction of phospho-CRTC3-S370 (Figure 6B). The TGFβ-induced cleavage of caspase-3 and PARP at 24 hours was blocked by SB-505124 (Figure 6C). Interestingly, in cells incubated with MRT199665 and TGFβ_1_, the appearance of cleaved caspase-3 and PARP were substantially enhanced compared with TGFβ-treated DMSO controls (Figure 6C). In the absence of TGFβ_1_, MRT199665 alone did not induce the cleavage of caspase-3 and PARP (Figure 6D). Furthermore, MRT199665+TGFβ_1_ treatment resulted in the maximal and more pronounced appearance of cleaved caspase-3 and PARP much earlier (12 hours) than TGFβ_1_ alone treatment (Figure 6D). When we monitored apoptosis using Annexin V and DAPI staining, treatment of NMuMG cells with TGFβ for 24 hours resulted in a substantial increase in Annexin V-positive apoptotic cells over controls, while this was reversed by SB-505124. Treatment of cells with MRT199665 significantly enhanced TGFβ-induced apoptosis (Figure 6E). Similarly, when we analysed cell viability, TGFβ_1_ treatment resulted in a decrease in the number of viable cells compared with DMSO control, while this was reversed by the SB-505124 (Figure 6F). In contrast, treatment of cells with MRT199665 resulted in almost complete loss of viable cells (Figure 6F). Collectively, these results indicate that inhibition of SIK isoforms by MRT199665 can potentiate TGFβ-mediated apoptotic cell death in NMuMG cells.

**Figure 6.**
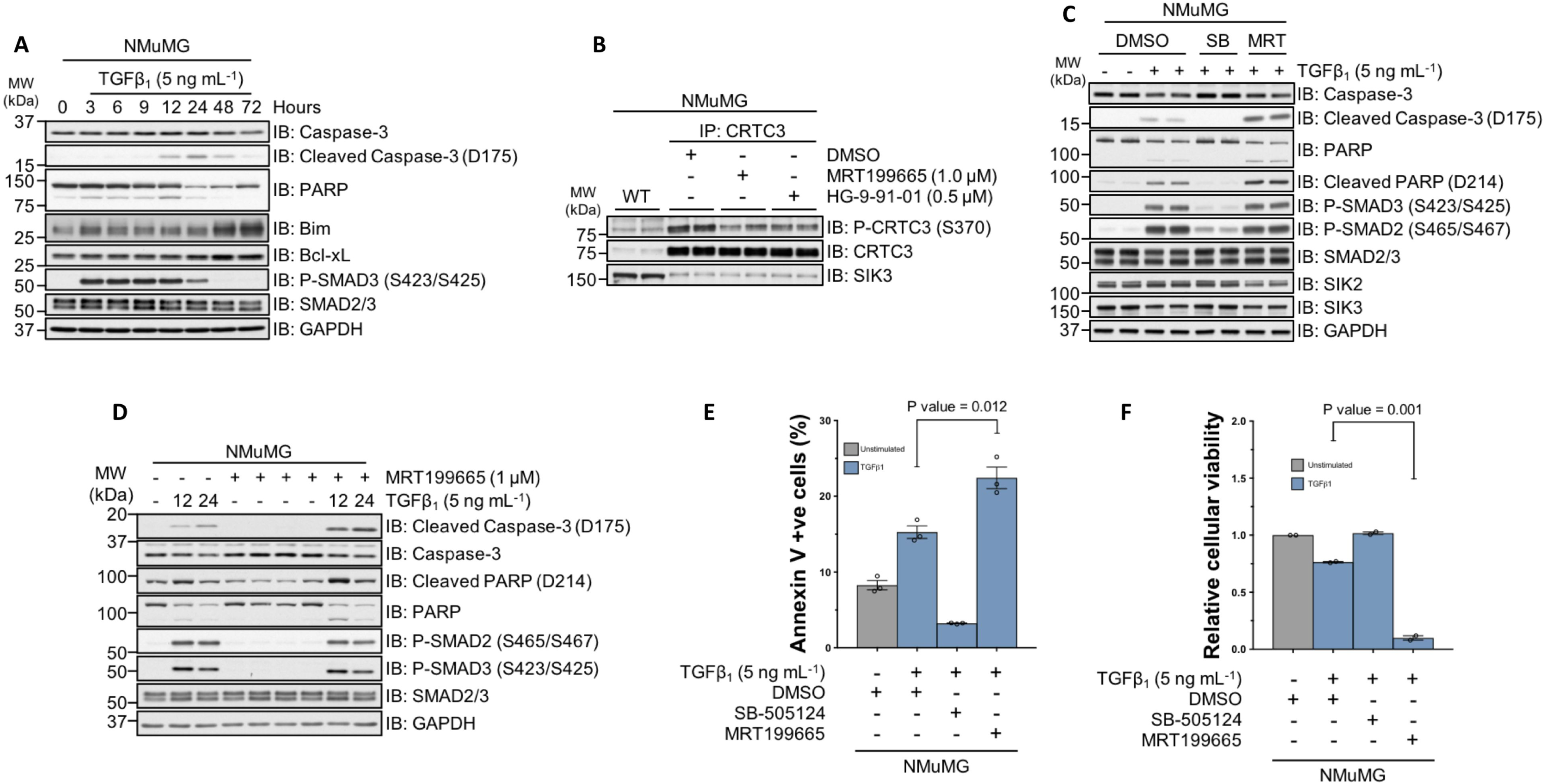
MRT199665 potentiates TGFβ-mediated apoptotic cell death. **A:** Immunoblot analysis of wild type NMuMG murine mammary epithelial cells stimulated with TGFβ_1_ for the indicated durations. Cell lysates were resolved via SDS-PAGE and membranes subjected to immunoblotting with the indicated antibodies. **B:** Immunoblot analysis of endogenous CRTC3 phosphorylation in wild type NMuMG cells following incubation with DMSO, MRT199665 or HG-9-91-01. Cell lysates were subjected to CRTC3 IP and subsequently resolved via SDS-PAGE. Membranes were subjected to immunoblotting with the indicated antibodies. **C:** Immunoblot analysis of wild type NMuMG cells incubated with either SB-505124 or MRT199665 in the presence of TGFβ_1_ stimulation. Cell lysates were resolved via SDS-PAGE and membranes subjected to immunoblotting with the indicated antibodies. **D:** Immunoblot analysis of wild type NMuMG cells incubated with MRT199665 in the presence or absence of TGFβ_1_ stimulation for the indicated durations. Cell lysates were resolved via SDS-PAGE and membranes subjected to immunoblotting with the indicated antibodies. **E:** Annexin V staining analysis of wild type NMuMG cells incubated with DMSO, SB-505124 or MRT199665 in the presence of TGFβ_1_ stimulation. Data represents the percentage of cells positive for annexin V staining from three independent experiments (5 x 10^4^ cell counts per sample per replicate). **F:** Crystal violet cellular viability analysis of wild type NMuMG cells incubated with DMSO, SB-505124 or MRT199665 in the presence of TGFβ_1_ stimulation. Data represents cellular viability relative to unstimulated DMSO control cells.

Previous reports have revealed that the clinically approved tyrosine kinase inhibitors (TKIs) bosutinib and dasatinib are also capable of inhibiting SIK isoforms, with *in vitro* IC_50_ values in the low nanomolar range^45, 46^ (Figure 7A). Both bosutinib and dasatinib inhibit the kinase activity of the BCR-Abl fusion as well as Src and BTK (Figure 7B) and are used to treat Philadelphia chromosome positive (Ph+) chronic myelogenous leukaemia (CML) and acute lymphoblastic leukaemia (ALL)^47, 48^. In WT U2OS cells, both compounds reduced the phospho-CRTC3 (S370) levels compared to DMSO controls, suggesting effective SIK inhibition (Figure 7C). Like MRT199665, neither bosutinib nor dasatinib inhibited the TGFβ-induced phosphorylation of SMAD2/SMAD3 but both inhibited the TGFβ-induced expression of GFP in U2OS 2G cells (Figure 7D), and endogenous PAI-1 in WT U2OS cells (Figure 7E). As in U2OS cells, in NMuMG cells, neither compound affected the TGFβ-induced phosphorylation of SMAD2/3 relative to controls (Figure 7F). Excitingly, treatment of NMuMG cells with bosutinib for 24 hours substantially enhanced the TGFβ-induced levels of cleaved caspase-3 and PARP to a similar extent as MRT199665 (Figure 7G), suggesting that the increased TGFβ-induced apoptosis caused by bosutinib is likely due to its ability to inhibit SIK isoforms.

**Figure 7.**
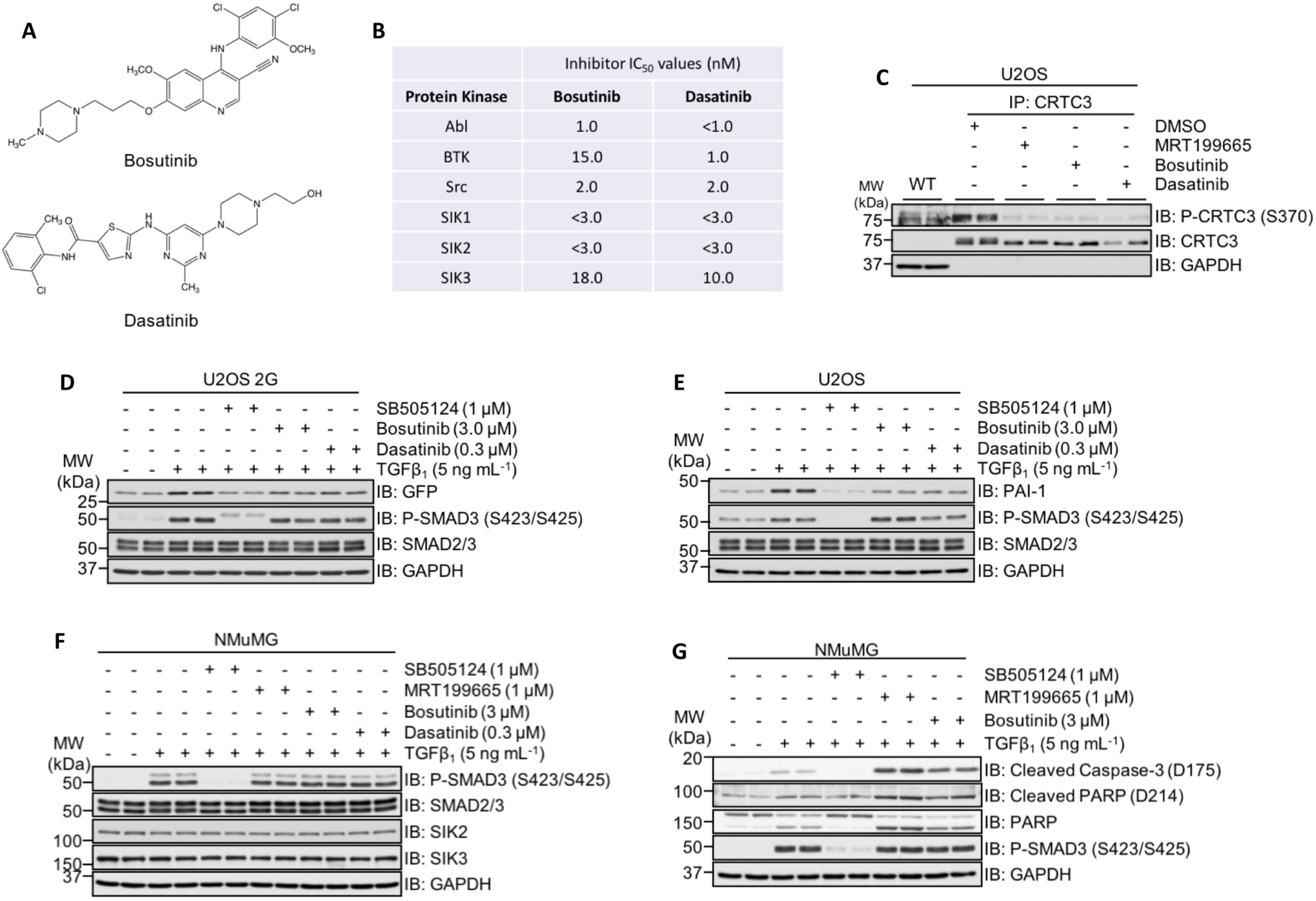
The clinically approved tyrosine kinase inhibitors (TKIs) bosutinib and dasatinib impact TGFβ-mediated transcriptional and cellular responses. **A:** The chemical structures of the clinically approved small-molecule tyrosine kinase inhibitors (TKIs) bosutinib and dasatinib. B: *In vitro* nanomolar IC_50_ values of bosutinib and dasatinib against the protein kinases Abl, BTK, Src and the three SIK isoforms (adapted from(Ozanne, Prescott and Clark, 2015)). **C:** Immunoblot analysis of endogenous CRTC3 phosphorylation in wild type U2OS cells following incubation with DMSO, MRT199665, bosutinib or dasatinib. Cell lysates were subjected to CRTC3 IP and subsequently resolved via SDS-PAGE. Membranes were subjected to immunoblotting with the indicated antibodies. **D:** Immunoblot analysis of U2OS 2G transcriptional reporter cells incubated with either SB-505124, bosutinib or dasatinib in the presence of TGFβ_1_ stimulation. Cell lysates were resolved via SDS-PAGE and membranes subjected to immunoblotting with the indicated antibodies. **E:** Immunoblot analysis of wild type U2OS cells incubated with either SB-505124, bosutinib or dasatinib in the presence of TGFβ_1_ stimulation. Cell lysates were resolved via SDS-PAGE and membranes subjected to immunoblotting with the indicated antibodies. **F:** Immunoblot analysis of wild type NMuMG cells incubated with SB-505124, MRT199665, bosutinib or dasatinib in the presence of TGFβ_1_ stimulation. Cell lysates were resolved via SDS-PAGE and membranes subjected to immunoblotting with the indicated antibodies. **G:** Immunoblot analysis of wild type NMuMG cells incubated with SB-505124, MRT199665 or bosutinib in the presence of TGFβ_1_ stimulation. Cell lysates were resolved via SDS-PAGE and membranes subjected to immunoblotting with the indicated antibodies.

### SIKs do not appear to affect SMAD2/3 phosphorylation and their nuclear translocation directly

We sought to investigate the molecular mechanisms by which SIK2/SIK3 might regulate TGFβ signalling. To explore whether SIKs exert effects on TGFβ signalling through direct phosphorylation of SMAD proteins, *in vitro* kinase assays were performed. GST-SIK2 and GST-SIK3, but not MBP-SIK1, phosphorylated SMAD2, SMAD3 and SMAD4 *in vitro* (Figure S1), while SIK inhibitor HG-9-91-01 blocked SMAD3 phosphorylation (Figure S2). Mass spectrometry identified Thr247 as the predominant SIK2/3 phosphorylated residue on SMAD3. This residue is conserved in SMAD2, SMAD3, SMAD4 and SMAD9 proteins (Figure 8A). The TGFβ-induced transcription of *PAI-1* has been previously reported to be specific to SMAD3^23, 49^. Consistent with this, treatment of wild type and SMAD3^-/-^ HaCaT cells with human TGFβ_1_ resulted in C-terminal phosphorylation of SMAD2 however PAI-1 expression was completely abrogated in SMAD3^-/-^ cells but not in WT cells (Figure 8B). Transient restoration of SMAD3 expression with FLAG-SMAD3^WT^ was sufficient to partially restore TGFβ-induced PAI-1 expression (Figure 8D). However, restoration of the SIK-phospho-deficient mutant FLAG-SMAD3^T247A^ in SMAD3^-/-^ cells also restored TGFβ-induced PAI-1 expression, to similar levels observed with SMAD3^WT^ (Figure 8B), suggesting that phosphorylation of SMAD3 at Thr247 by SIKs is unlikely to explain effects of SIKs in TGFβ signalling.

**Figure 8.**
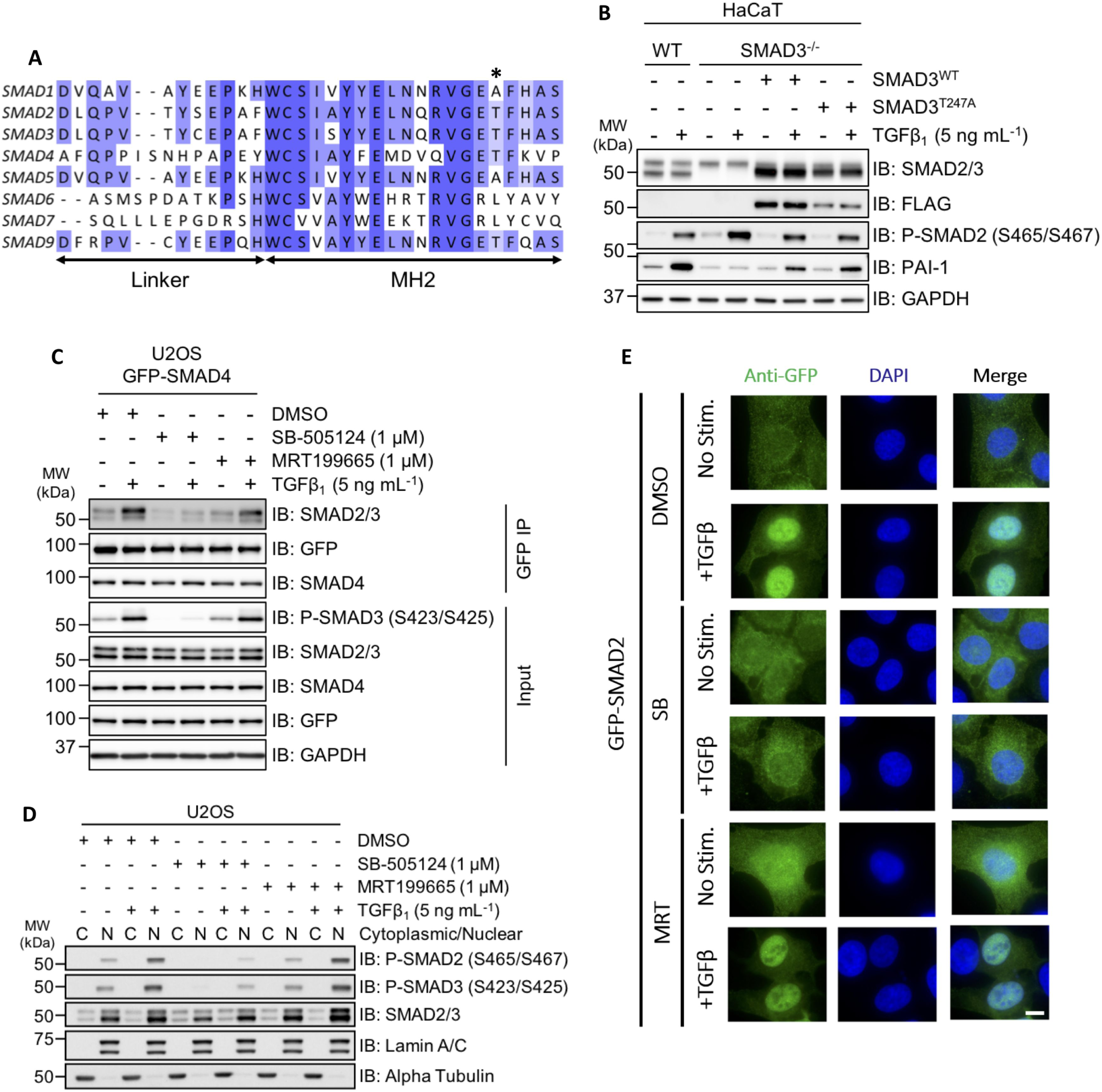
SIK2 and SIK3 isoforms phosphorylate SMAD3 *in vitro* however MRT199665 does not affect SMAD interaction or nuclear translocation. A: Sequence alignment of the eight human SMAD proteins. The asterisk indicates the amino acid residue in recombinant SMAD3 (Thr247) that was identified via phosphorylation-site mapping analysis. **B:** Immunoblot analysis of wild type HaCaT cells and SMAD3^-/-^ cells following transient overexpression of either FLAG-SMAD3^WT^ or FLAG-SMAD3^T247A^ mutant, in the presence or absence of TGFβ_1_ stimulation. Cell lysates were resolved via SDS-PAGE and membranes subjected to immunoblotting with the indicated antibodies. **C:** U2OS cells stably expressing GFP-SMAD4 were incubated with DMSO, SB-505124 or MRT199665 in the presence or absence of TGFβ_1_ stimulation. Cell lysates were subjected to GFP immunoprecipitation and resolved via SDS-PAGE. Membranes were subjected to immunoblotting with the indicated antibodies. **D:** Immunoblot analysis of SMAD2 and SMAD3 localisation in wild type U2OS cells following incubation with either DMSO, SB-505124 or MRT199665, in the presence or absence of TGFβ_1_ stimulation. Cell lysates were separated into cytoplasmic (C) and nuclear (N) fractions, resolved via SDS-PAGE and membranes subjected to immunoblotting with the indicated antibodies. **E:** Immunofluorescence analysis of U2OS cells stably expressing GFP-SMAD2 following incubation with DMSO, SB-505124 or MRT199665 in the presence or absence of TGFβ_1_ stimulation (images are representative). Scale bar indicates 10 µm.

Next, we whether SIK2/SIK3 inhibition disrupts the formation of SMAD2/SMAD3-SMAD4 complexes or nuclear accumulation of SMADs. In U2OS cells stably overexpressing GFP-SMAD4, neither the basal nor TGFβ-induced increase in co-precipitation of SMAD2/3 in GFP-SMAD4 IPs was affected by treatment of cells with MRT199665 (Figure 8C). TGFβ-induced a robust nuclear accumulation of phosphorylated SMAD2 and SMAD3 levels in nuclear fractions over unstimulated conditions or when cells were treated with SB-505124 (Figure 8D). MRT199665 treatment did not affect the cytoplasmic/nuclear distribution of phosphorylated SMAD2/SMAD3 relative to controls in both unstimulated and TGFβ_1_ stimulated conditions (Figure 8D). Consistent with this, in cells stably overexpressing GFP-SMAD2, MRT199665 did not prevent the nuclear translocation of GFP-SMAD2 following TGFβ_1_ stimulation when analysed via immunofluorescence (IF) (Figure 8E). As inhibition of SIK isoforms with MRT199665 does not appear to impact the formation of the SMAD2/3-SMAD4 complex or the nuclear accumulation of phosphorylated SMAD2/3, the effect of MRT199665 is likely to occur further downstream.

## Discussion

In this study, we identified inhibitors of SIK isoforms as novel candidates for the inhibition of TGFβ-induced transcription. We established that inhibiting SIK protein kinase activity, both pharmacologically and genetically, attenuates the TGFβ-induced expression of endogenous *PAI-1* transcript and protein in different cells. Moreover, the attenuation of TGFβ-induced PAI-1 by MRT199665 occurred without affecting phosphorylation of SMAD proteins, the SMAD2/SMAD3-SMAD4 interaction or the nuclear accumulation of activated SMADs. We propose that SIK isoforms function at the level of transcriptional regulation in the context of TGFβ signalling.

In every cell line we tested, TGFβ-induced endogenous PAI-1 transcript and protein levels were reduced by both pharmacological and genetic ablation of SIK kinase activity. PAI-1 is a serine protease inhibitor (Serpin) that functions as the physiological inhibitor of the serine proteases tissue-type plasminogen activator (t-PA) and urokinase-type plasminogen activator (u-PA) and controls fibrinolysis. Increased plasma levels of PAI-1 have been associated with a number of diseases including thrombotic vascular disorders^50^. As inhibition of SIK isoforms attenuates TGFβ-induced expression of PAI-1, pharmacological SIK inhibitors may have novel therapeutic potential if they demonstrably suppress excessive PAI-1 levels *in vivo*.

SIK1 was previously linked to the regulation of TGFβ signalling^51, 52^ in which it was shown that SIK1 was a direct transcriptional target of TGFβ signalling and played a role in the degradation of ALK5 through SIK1/SMAD7/SMURF2 complex. During the course of our experiments, we did not observe any change in protein of expression of either SIK2 or SIK3 upon TGFβ signalling, suggesting that unlike SIK1, these two isoforms are not transcriptional targets of TGFβ signalling. Our data indicates that the inhibition of the kinase activity of SIK2 and SIK3 is sufficient to suppress the TGFβ-induced upregulation of PAI-1, whereas RNAi-mediated depletion of SIK1 has been reported to enhance *PAI-1* mRNA expression in response to TGFβ stimulation^51^. It is therefore apparent that the exact roles of the different SIK isoforms in regulating TGFβ signalling remains to be elucidated and is most likely context dependent.

The precise mechanisms by which SIK isoforms regulate the TGFβ-induced expression of *PAI-1* or other genes remains to be resolved. As serine-threonine protein kinases, SIK isoforms act by phosphorylating protein substrates. In the case of TGFβ signalling, these could be components of the SMAD-transcriptional complexes or key transcriptional modulators, enhancers, suppressors as and/or adaptors that modulate the function of these transcriptional co-factors in order to control the transcriptional activity of SMAD2/SMAD3. Unless the core SMAD2/3 transcriptional complexes are found to be substrates of SIKs, the impact of SIKs in TGFβ-target gene transcription is likely to be determined by whether the individual target gene promoters recruit specific SIK substrates. It is known that SMAD2/SMAD3 do not directly regulate target gene transcription, but instead facilitate the recruitment of various transcriptional co-activators/co-repressors or histone modifying enzymes^20^. SIKs have been reported to regulate the Toll-like Receptor (TLR) signalling through their ability to phosphorylate the transcriptional coactivator CRTC3 and so reduce CREB-dependent transcription of the *IL10* gene^33^. It is therefore conceivable that SIKs may employ similar mechanisms to regulate SMAD-associated transcriptional co-factors to modulate the transcription of specific subsets of TGFβ-target genes. A phospho-proteomic approach using both SIK inhibitors and SIK2^T175A^/SIK3^T163A^ MEFs might uncover potential SIK substrates that underpin the regulation of TGFβ-induced transcription of distinct genes.

In many epithelial cell types, TGFβ-induced cytostasis through p15^INK4B^, p21^CIP1^ and p27^KIP1^ ^1,4^ is often followed by context-dependent cell fates, such as differentiation, epithelial-mesenchymal transition (EMT) or apoptosis^53, 54^. Interestingly, inhibition of SIKs induced p21^CIP1^ and p27^KIP1^ expression independently of TGFβ stimulation. Perhaps due in part to this, SIK inhibitors sensitise NMuMG cells for TGFβ-induced apoptosis. These observations imply that SIK inhibitors could be employed to sensitise cancer cells for apoptosis in those cells that TGFβ induces apoptosis. This could be easily tested by using clinically approved anti-cancer drugs dasatinib and bosutinib that inhibit SIK isoforms in addition to their intended targets in a number of cancer cell types, including multiple Burkitt’s lymphoma (BL) cell lines^42, 55–57^, hepatocellular carcinoma^58^ and prostate carcinoma cells^59–61^ that have been reported to undergo apoptosis in response to TGFβ.

Our findings place SIK isoforms as modulators of a subset of TGFβ-induced transcriptional and physiological responses. Understanding these in detail will allow targeting of selective TGFβ responses, thereby limiting potential consequences of inhibiting the TGFβ pathway in its entirety. Of course, as discussed above, SIKs themselves are known to control other pathways, including immune signalling, and these need to be considered carefully. Taking into consideration that the clinically approved TKIs dasatinib and bosutinib, which also potently inhibit SIK isoforms^45, 46^, are administered to patients safely, it is conceivable that more specific SIK inhibitors could be applied to target certain TGFβ-associated pathologies.

## Materials and methods

### Antibodies

For Western immunoblotting analysis, all primary IgG antibodies were used at 1:1000 dilution unless otherwise stated. Anti-Phospho-SMAD3 (S423/S425) Rabbit polyclonal IgG (600-401-919) was purchased from Rockland Inc. Anti-GFP Mouse monoclonal IgG (11814460001) was purchased from Roche. Anti-Phospho-SMAD2 (S465/S467) Rabbit polyclonal IgG (3101), anti-SMAD2/3 Rabbit monoclonal IgG (8685), anti-c-Myc Rabbit monoclonal IgG (5605), anti-p27^KIP1^ Rabbit monoclonal IgG (3688), anti-p21^WAF1/CIP1^ Rabbit monoclonal IgG (2947), anti-GAPDH Rabbit monoclonal IgG (used at 1:5000 dilution) (2118), anti-SIK2 Rabbit IgG (6919) were all purchased from Cell Signalling Technology (CST). Anti-PAI-1 Rabbit polyclonal IgG (ab66705), anti-CTGF Rabbit polyclonal IgG (ab6992) and anti-CRTC3 Rabbit monoclonal IgG (ab91654) were purchased from Abcam. Anti-Phospho-CRTC3 (S370) Sheep polyclonal IgG (S253D, 3^rd^ bleed) and anti-SIK3 Sheep polyclonal IgG (S373D, 3^rd^ bleed) were generated by MRC PPU Reagents and Services. Species-specific horseradish peroxidase (HRP)-conjugated secondary antibodies were used at 1:2500-5000 dilution. Rabbit anti-Sheep polyclonal IgG (H+L) Secondary Antibody, HRP (31480) and Goat anti-Mouse polyclonal IgG (H+L) Secondary Antibody, HRP (31430) were purchased from Thermo Fisher Scientific. Goat anti-Rabbit polyclonal IgG (H+L), HRP-conjugated Secondary Antibody (7074) was purchased from CST.

### Cytokines and pharmacological inhibitors

Purified recombinant human TGFβ_1_ was purchased from either R&D Systems or PeproTech and reconstituted in sterile 4 mM HCl containing 1 mg mL^-1^ bovine serum albumin (BSA). Prior to stimulation with TGFβ_1_, cells were cultured in serum-free culture media for approximately 16 hours at 37°C in order to reduce autocrine signalling. Pharmacological inhibitors were reconstituted at 10 mM in dimethyl sulfoxide (DMSO) and used at the concentrations and durations indicated in the respective figure/figure legend. For all inhibitor experiments, control cells were incubated with an equivalent volume of DMSO.

### Generation of SIK KI mice

SIK2^T175A^/SIK3^T163A^ homozygous kinase dead knock-in (KI) mice were bred from SIK2^tm1.1Arte^ and SIK3^tm1.1Arte^ mice maintained on a C57BL/6NJ genetic background as described previously^39^. Primary mouse embryonic fibroblasts (MEFs) were generated from SIK2^T175A^/SIK3^T163A^ or SIK3^WT^ embryos at E11.5-13.5 as detailed previously^62^. Primary and SV-40 immortalised MEFs were cultured in DMEM supplemented with 20% (v/v) FBS, 2 mM *L*-glutamine, 100 units mL^-1^ Penicillin, 100 µg mL^-1^ Streptomycin, 1x Minimum Essential Medium (MEM) Non-Essential Amino Acids (NEAA) and 1 mM Sodium Pyruvate. Mice were maintained in individually ventilated cages and provided with free access to food and water under specific pathogen-free conditions consistent with E.U. and U.K. regulations. All animal breeding and studies were conducted following approval by the University of Dundee Ethical Review Committee and performed under a U.K. Home Office Project Licence granted under the Animals (Scientific Procedures) Act 1986.

### Mammalian cell culture

A-172 human glioblastoma, A549 human pulmonary adenocarcinoma, U2OS human osteosarcoma, HACAT human immortalised keratinocyte, HEK-293 human embryonic kidney and HeLa human cervical adenocarcinoma cells were obtained from the MRC PPU Tissue Culture facility and cultured in Dulbecco’s Modified Eagle’s Medium (DMEM) supplemented with 10% (v/v) fetal bovine serum (FBS), 2 mM *L*-glutamine, 100 units mL^-1^ Penicillin and 100 µg mL^-1^ Streptomycin (hereafter referred to as D10F media). NMuMG murine mammary epithelial cells were cultured in D10F media supplemented with 10 µg mL^-1^ insulin (from bovine pancreas). All cell lines were maintained at 37°C in a humidified atmosphere with 5% (v/v) CO_2_ levels and routinely tested for mycoplasma contamination.

### Generation of *SMAD3^-/-^* knockout cells using CRISPR-Cas9

To generate *SMAD3^-/-^* knockout cells by CRISPR-Cas9 genome editing, HaCaT cells were transfected with the plasmid pSpCas9(BB)-2A-GFP (PX458)^63^ containing both the Cas9 endonuclease and a guide RNA (gRNA) pair targeting exon 6 of the endogenous *SMAD3* gene. For the acquisition of single-cell knockout clones, single cells were isolated by fluorescence-activated cell sorting (FACS) and plated in individual wells of 96-well cell culture plates. Viable cell clones were expanded, and successful knockouts confirmed by both Western immunoblotting and genomic DNA sequencing. Sequence of gRNA oligonucleotides: SMAD3 forward gRNA 5’-CACCGGAATGTCTCCCCGACGCGC-3’; SMAD3 reverse gRNA 5’-AAACGCGCGTCGGGGAGACATTCC-3’.

### Mammalian cell lysis

Cells were washed twice with cold 1x DPBS and incubated with lysis buffer (50 mM Tris/HCl pH 7.5, 270 mM sucrose, 150 mM sodium chloride, 1 mM EDTA pH 8.0, 1 mM EGTA pH 8.0, 1 mM sodium orthovanadate, 10 mM sodium β-glycerophosphate, 50 mM sodium fluoride, 5 mM sodium pyrophosphate, 1% (v/v) Nonidet P-40 (NP-40)) supplemented with Complete, EDTA-free Protease Inhibitors (Roche) (one tablet per 25 mL) for approximately 5 minutes on ice. Cell lysates were transferred to 1.5 mL microcentrifuge tubes and centrifuged at 16,000 x g for 10 minutes at 4°C and either processed immediately or cryopreserved in liquid nitrogen prior to storage at −80°C. The protein concentrations of the cell lysate samples were determined using Pierce Coomassie (Bradford) Protein Assay Kit (Thermo Fisher Scientific). Cell lysate samples were subsequently diluted using 4x NuPAGE LDS (lithium dodecyl sulfate) sample buffer (Invitrogen) supplemented with 8% (v/v) 2-Mercaptoethanol (2-ME) (Sigma-Aldrich) and the sample concentrations equalised.

### Luciferase transcriptional reporter assay

U2OS 2G transcriptional reporter cells were seeded in 6-well cell culture plates and incubated with the required small-molecule inhibitors/cytokine at the indicated concentrations and duration. Cells were subsequently washed twice with 1x DPBS and lysed using 1x Cell Culture Lysis Reagent (CCLR; Promega). Cell culture plates were incubated for approximately 5 minutes on bench-top platform rocker to ensure efficient cell lysis. Cell lysates were transferred to 1.5 mL microcentrifuge tubes and kept on ice. Lysate samples were vortexed for approximately 10 seconds, centrifuged at 12,000 x g for 2 minutes at 4°C and 200 µL of supernatant transferred to new 1.5 mL microcentrifuge tubes. Lysate samples were subsequently transferred to 96-well white flat-bottom cell culture microplates. An equivalent volume of 2x Luciferase Assay Buffer (50 mM Tris/Phosphate pH 7.8, 16 mM MgCl_2_, 2 mM dithiothreitol (DTT), 1 mM adenosine triphosphate (ATP), 30% (w/v) glycerol, 1% (w/v) bovine serum albumin (BSA), 250 µM *D*-Luciferin, 8 µM sodium pyrophosphate) was subsequently added to each well and the microplate incubated for approximately 1 minute on a bench-top vibrating platform. Luminescence values were obtained using an EnVision 2104 Multimode Microplate Reader (PerkinElmer). The protein concentrations of each lysate sample were determined using Pierce Coomassie (Bradford) Protein Assay Kit (Thermo Fisher Scientific) and used to normalise luminescence values.

### Cellular fractionation

Extraction of separate cytoplasmic and nuclear protein fractions from cultured U2OS cells was performed using NE-PER Nuclear and Cytoplasmic Extraction Reagents (Thermo Fisher Scientific) according to the manufacturer’s protocol. The supplied lysis buffers (CER I and NER) were supplemented with 1x Complete, EDTA-free Protease Inhibitors (Roche) immediately prior to use. Subcellular fractions were reduced using NuPAGE 4x LDS sample buffer containing 8% (v/v) 2-mercaptoethanol and incubated at 95°C for 5 minutes prior to SDS-PAGE. Fractions were resolved by SDS-PAGE and analysed via Western immunoblotting.

### Quantitative reverse transcription polymerase chain reaction (RT-qPCR)

For all RT-qPCR experiments, cells were seeded in 6-well cell culture plates and incubated with the required TGFβ_1_/inhibitor combinations for the durations indicated in the respective figure legends. Total RNA was isolated from the cells using the RNeasy Micro Kit (Qiagen) according to the manufacturer’s protocol. Complementary DNA (cDNA) was synthesised from 0.5-1.0 µg of isolated RNA using iScript cDNA Synthesis Kit (Bio-Rad) according to the manufacturer’s protocol. All RT-qPCR reactions were conducted in triplicate and included

50% (v/v) SsoFast EvaGreen Supermix (Bio-Rad), 0.5 µM forward primer, 0.5 µM reverse primer and the required volume of cDNA. RT-qPCR experiments were performed using CFX96 or CFX384 Real-Time PCR Detection Systems (Bio-Rad). The Ct (cycle threshold) values for each gene of interest were normalised to the arithmetic mean Ct value of the reference gene glyceraldehyde-3-phosphate dehydrogenase (GAPDH) or 18S ribosomal RNA (rRNA) using Microsoft Excel software. The 2^−ÄÄCt^ relative quantification method was then used to analyse the relative changes in gene expression between control and treatment conditions^64^. GraphPad Prism software (version 8.0) was used to generate graphs and perform statistical analysis.

### Immunofluorescence (IF) microscopy

U2OS GFP-SMAD2 cells were plated onto glass coverslips and treated as described in the respective figure legends. Cells were fixed in 4% (w/v) paraformaldehyde (PFA) immediately after aspirating culture media for 15 minutes at RT before washing twice in 1x PBS. Permeabilisation was performed using 0.2% Triton X (Sigma) in PBS for 10 minutes at RT before washing twice more in 1x PBS and blocking for 1 hour at RT with 5% (w/v) BSA in PBS. Primary antibody (anti-GFP polyclonal IgG, 1:1000 dilution) in 0.5% BSA/0.2% TWEEN 20 (Sigma)/PBS was added to coverslips for 1 hour at 37°C before washing three times in 0.5% BSA/0.2% TWEEN 20/PBS (10 minutes per wash). Cells were incubated with the secondary Alexa-Fluor conjugated antibody (anti-Rabbit, 488 nm; 1:500 dilution) for 1 hour at RT, before washing with 0.5% BSA/0.2% TWEEN 20 /PBS three times (20 minutes per wash). During the second wash of the three, 4’,6-Diamidino-2-Phenylindole Dihydrochloride (DAPI; Sigma) was added at a final concentration of 1 µg mL^-1^ and removed in the final wash. Coverslips were rinsed in deionised H_2_O and mounted onto glass microscopy slides using VECTASHIELD (Vector Laboratories). Coverslips were sealed and left to dry overnight at 4°C. Cells were imaged using the DeltaVision Imaging System (20x or 60x objective; GE Healthcare) and processed using softWoRx (GE Healthcare) and OMERO^65^.

### Annexin V staining assay

NMuMG cells were incubated with the required cytokine/inhibitor combinations. Following this, both adherent and non-adherent (*i.e.* apoptotic) cells were collected into 50 mL conical centrifuge tubes, pelleted by centrifugation (300 x g, 2 minutes) and washed once using cold 1x DPBS. Cells were subsequently centrifuged (300 x g, 2 minutes), the cell pellets were resuspended in Annexin Binding Buffer, ABB (10 mM HEPES, 140 mM NaCl, 2.5 mM CaCl_2_, pH 7.4) and transferred to 1.5 mL microcentrifuge tubes. The required cell suspension samples were then incubated with Annexin V, Alexa Fluor 488 conjugate (Invitrogen; A13201) for 15 minutes at RT and protected from light. The appropriate samples were subsequently incubated with 5 µg mL^-1^ DAPI (4’,6-diamidino-2-phenylindole) (Invitrogen). Samples were immediately analysed using a BD LSRFortessa Cell Analyser (BD Biosciences) and BD FACSDiva acquisition software (BD Biosciences). Annexin V Alexa Fluor 488 fluorescence was detected by excitation at 488 nm and emission at 530 ± 30 nm, and DAPI fluorescence was detected by excitation at 355 nm and emission at 450 ± 50 nm. Single cells were identified on the basis of forward light scatter (FSC) and side light scatter (SSC), and subsequently evaluated for Annexin V Alexa Fluor 488 and DAPI fluorescence. Data analysis was performed using FlowJo Single Cell Analysis Software (BD Biosciences).

### Cellular proliferation

HaCaT cells were assayed for proliferation using an IncuCyte ZOOM Live-Cell Analysis System (Sartorius). Cells were plated in 96-well cell culture plates (1 x 10^3^ cells per well, 6 wells per condition, plates in triplicate) and incubated with the required small-molecule inhibitor in the presence or absence of recombinant human TGFβ_1_ (5 ng mL^-1^). Cells were imaged every 2 hours over a period of 170 hours and the percent confluence determined using IncuCyte ZOOM Analysis Software (Sartorius).

### Crystal violet cellular viability assay

NMuMG cells were seeded in 96-well cell culture plates and incubated for 24 hours at 37°C to enable adherence of cells to culture plates. The inclusion of wells containing culture medium without cells were used as negative control wells. Following initial 24-hour incubation, culture media was aspirated and replaced with reduced serum (1% v/v FBS) DMEM supplemented with 10 µg mL^-1^ bovine insulin (Sigma-Aldrich) containing the required inhibitors or equivalent volume DMSO, with or without recombinant human TGFβ_1_ (5 ng mL^-1^) and incubated for a further 24 hours. Cells were subsequently fixed using 10% (v/v) methanol/10% (v/v) acetic acid for 5 minutes at RT and subsequently washed with 1x PBS. Fixed cells were stained using 0.5% (w/v) crystal violet staining solution (0.5 g crystal violet powder (Sigma-Aldrich), 80 mL distilled H_2_O and 20 mL methanol) for 20 minutes at RT on a bench-top platform rocker. Plates were subsequently washed carefully using tap water, inverted on filter paper to remove residual liquid and allowed to air-dry overnight. Following this, methanol was added to each well and incubated for 20 minutes at RT on a bench-top platform rocker. The absorbance value of each well was measured at 570 nm (OD_570_) using a 96-well microplate spectrophotometer. The mean OD_570_ value of negative control wells (*i.e.* wells not containing cells) was subtracted from the values obtained from each well on the culture plate and the percentage of viable cells for each condition determined relative to the mean average OD_570_ value of non-stimulated DMSO control treated cells.

### *In vitro* protein kinase assay

21 µL reaction solutions were prepared containing 200 ng of protein kinase and 2 µg of substrate protein in 1x kinase assay buffer (50 mM Tris-HCl pH 7.5, 0.1 mM EGTA, 10 mM magnesium acetate, 0.1% (v/v) 2-mercaptoethanol and 0.1 mM [γ^32^P]-ATP). Reactions were conducted at 30°C for 30 minutes at 1050 rpm and terminated via the addition of 7 µL NuPAGE 4x LDS sample buffer containing 8% (v/v) 2-mercaptoethanol. For *in vitro* kinase assays involving the use of small-molecule inhibitors, reaction solutions containing all the required components were incubated at 30°C for 10 minutes at 1050 rpm prior to the addition of 0.1 mM [γ32P]-ATP. Reactions were then performed as detailed previously. Samples were incubated at 95°C for 5 minutes and subsequently centrifuged at 5.0 x 10^3^ x g for 1 minute. Samples were loaded onto NuPAGE 4-12% Bis-Tris precast polyacrylamide gels and resolved via SDS-PAGE. Polyacrylamide gels were subsequently stained with InstantBlue Coomassie Protein Stain (Expedeon) to visualise the resolved recombinant proteins and imaged using the ChemiDoc Imaging System (Bio-Rad). ^32^P radioactivity was analysed via autoradiography using Amersham Hyperfilm (GE Healthcare Life Sciences).

### Statistical analysis

All experiments have a minimum of three biological replicates unless otherwise stated in the respective figure legend. In addition, all luciferase, RT-qPCR, cellular proliferation, annexin V staining and crystal violet staining experiments have at least three technical repeats for each biological replicate. The data are presented as the arithmetic mean with error bars denoting the standard error of the mean (SEM). The statistical significance of differences between experimental conditions were assessed using either Student’s t-test or analysis of variance (ANOVA) with Bonferroni correction using GraphPad Prism (version 8.0) analysis software. Differences in the mean of experimental conditions was considered significant if the probability value (p-value) was <0.05. All immunoblotting figures are representative.

## Acknowledgments

We thank staff at the MRC PPU International Centre for Kinase Profiling (University of Dundee, UK) for providing us with the inhibitor panel used for screening. We thank E. Allen, J. Stark, and A. Muir for help and assistance with tissue culture, and the cloning, antibody and protein production teams within MRC PPU Reagents and Services (University of Dundee, UK), coordinated by J. Hastie and H. McLauchlan. We thank Dr. R. Clarke from the flow cytometry facility (School of Life Sciences, University of Dundee, UK) for her invaluable help and advice with Annexin V staining assays.

## Supplementary Figures and Legends

**Figure S1.**
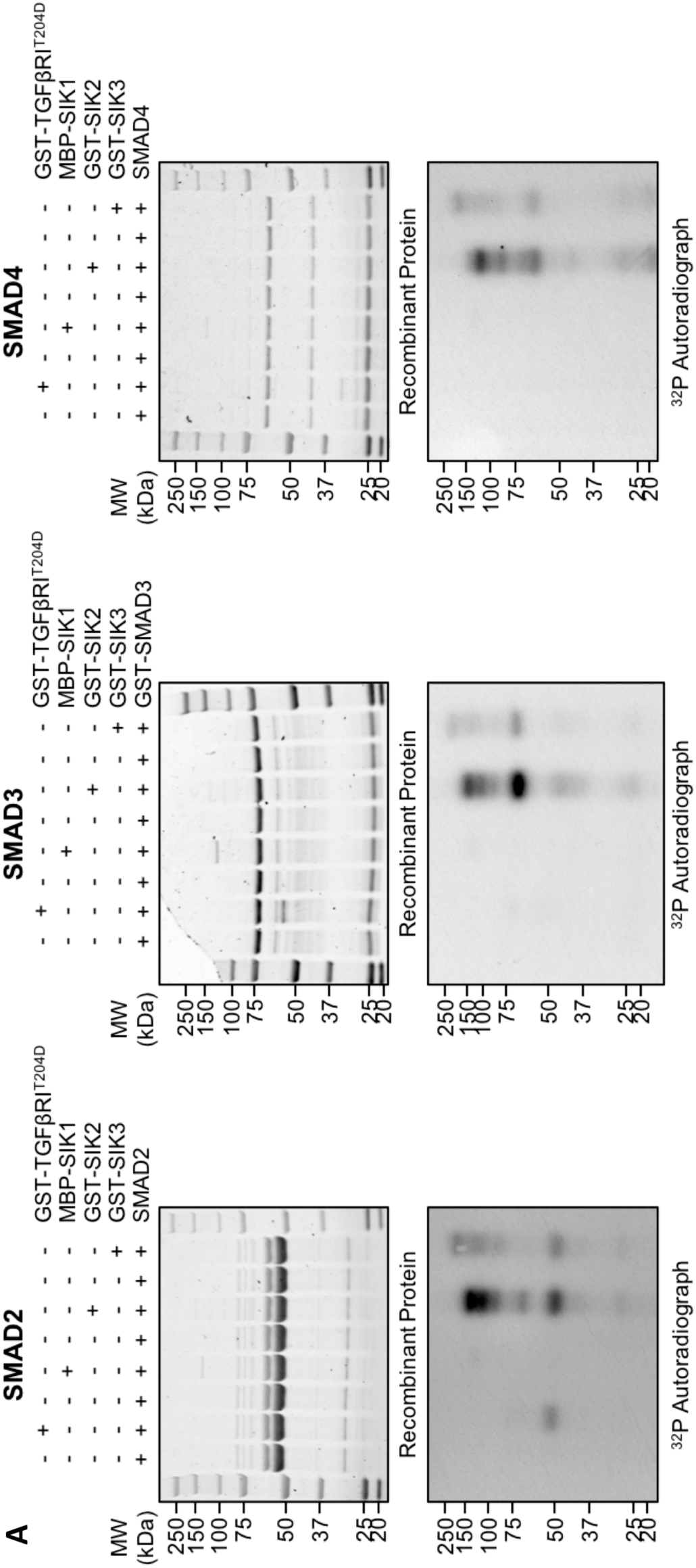
*In vitro* phosphorylation of recombinant SMAD proteins. **A:** *In vitro* protein kinase assay analysis using the recombinant kinases MBP-SIK1, GST-SIK2 and GST-SIK3 and either recombinant SMAD2, SMAD3 or SMAD4 as the substrate. A constitutively active mutant form of the type I TGFβ receptor (GST-TGFβRIT204D) was included as a positive control.

**Figure S2.**
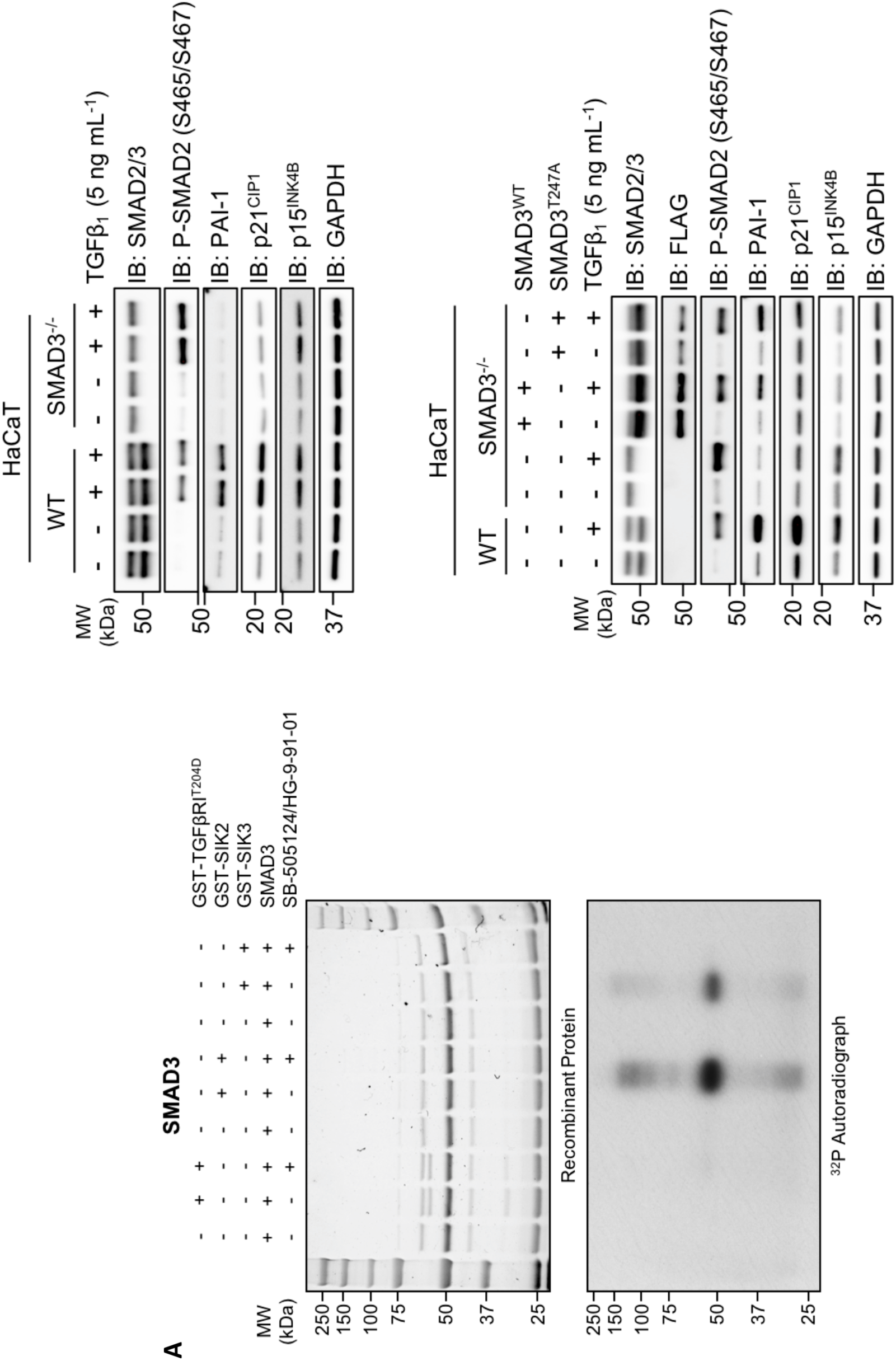
**A:** *In vitro* protein kinase assay analysis using the recombinant kinases GST-TGFβRIT204D, GST-SIK2 and GST-SIK3 and cleaved recombinant SMAD3 as the substrate. The small-molecule inhibitors SB-505124 and HG-9-91-01 were included as controls. **B:** Immunoblot analysis of wild type HaCaT cells and SMAD3-/- cells following incubation with recombinant human TGFβ1. Cell lysates were resolved via SDS-PAGE and membranes subjected to immunoblotting with the indicated antibodies. **C:** Immunoblot analysis of wild type HaCaT cells and SMAD3-/- cells following transient overexpression of either FLAG-SMAD3WT or FLAG-SMAD3T247A mutant, in the presence or absence of TGFβ1 stimulation. Cell lysates were resolved via SDS-PAGE and membranes subjected to immunoblotting with the indicated antibodies.

